# An expanded GCaMP reporter toolkit for functional imaging in *C. elegans*

**DOI:** 10.1101/2023.03.06.531342

**Authors:** Jimmy Ding, Lucinda Peng, Sihoon Moon, Hyun Jee Lee, Dhaval S. Patel, Hang Lu

## Abstract

In living organisms, changes in calcium flux are integral to many different cellular functions and are especially critical for the activity of neurons and myocytes. Genetically encoded calcium indicators (GECIs) have been popular tools for reporting changes in calcium levels *in vivo*. In particular, GCaMP, derived from GFP, are the most widely used GECIs and have become an invaluable toolkit for neurophysiological studies. Recently, new variants of GCaMP, which offer a greater variety of temporal dynamics and improved brightness, have been developed. However, these variants are not readily available to the *Caenorhabditis elegans* research community. This work reports a set of GCaMP6 and jGCaMP7 reporters optimized for *C. elegans* studies. Our toolkit provides reporters with improved dynamic range, varied kinetics, and targeted subcellular localizations. Besides optimized routine uses, this set of reporters are also well-suited for studies requiring fast imaging speeds and low magnification or low-cost platforms.

## Introduction

Calcium ions play a significant role in many biological processes, such as apoptosis, transcription, muscle contraction, and neuronal excitability (CLAPHAM 2007). Imaging intracellular calcium flux using GECIs is an invaluable tool for *in vivo* measurement of neural activity. Continual advances in GECIs have resulted in brighter, more sensitive reagents with faster kinetics, such as the GCaMP6 (CHEN *et al*. 2013) and jGCaMP7 (DANA *et al*. 2019) families of fluorophores. These higher signal-to-noise fluorophores and advances in optical microscopy have allowed more reliable measurement of neural activity during demanding experiments, such as recording from many neurons in parallel (AHRENS *et al*. 2013; KATO *et al*. 2015; LI *et al*. 2017; WIRAK *et al*. 2022), neural processes (HENDRICKS *et al*. 2012; PARKER *et al*. 2016; LIU *et al*. 2018), and whole-brain imaging in freely behaving animals (NGUYEN *et al*. 2016; VENKATACHALAM *et al*. 2016; KIM *et al*. 2017; ZONG *et al*. 2022).

In the model organism *C. elegans*, GCaMP has been widely used to determine the roles of individual neurons within the nervous system. For example, the expression of cytosolic GCaMP in different classes of neurons has mapped their activities to specific stimuli (ZASLAVER *et al*. 2015), mapped circuits that drive olfactory behaviors (CHALASANI *et al*. 2007; HA *et al*. 2010), shown experience-dependent changes in neural activity (HA *et al*. 2010; HONG *et al*. 2017; WU *et al*. 2023), and verified the roles of key hub neurons in sustaining behavioral states (FLAVELL *et al*. 2013) among many applications. Additionally, it has been shown that subcellular (HENDRICKS *et al*. 2012) and multicellular (GORDUS *et al*. 2015; KATO *et al*. 2015) calcium activity play important roles in the neurophysiology of *C. elegans*. Such studies impose greater performance demands on optical microscopy systems as they confine fluorophores to specific cell regions, such as nuclei for multicell imaging and processes for subcellular imaging, which comes with reduced signal (or signal-to-noise ratio). The newer generation GCaMP reagents (GCaMP6 and jGCaMP7) (CHEN *et al*. 2013; DANA *et al*. 2019) can alleviate some of these technical challenges. However, *C. elegans*-optimized versions of GCaMP6f, jGCaMP7s, and 7f remain publicly unavailable, and the *in vivo* properties of jGCaMP7 variants have not been fully characterized in *C. elegans*.

To address the *C. elegans* community’s need, we have made a toolkit of codon-optimized reagents that includes four variants with distinct kinetic profiles (GCaMP6f, 6s, jGCaMP7f, 7s), each available with three subcellular localizations - cytosolic, nuclear, and membrane-targeted. The kinetic variants offer flexibility in balancing brightness and speed for different experiments. At the same time, the subcellular localization tags enable imaging of calcium flux in specific parts of a neuron. For example, membrane localization allows neurite recording, while nuclear localization enables the separation of signals from multiple neurons recorded in parallel. Additionally, to facilitate the detection of neurons and neurites during fluctuations in GCaMP brightness and reduce motion artifacts when processing data, our reporters are bicistronic and express the bright red fluorophore mScarlet-I (BINDELS *et al*. 2017) in tandem with each GCaMP variant.

To compare and contrast the properties of each GCaMP variant *in vivo*, we generated single-copy knock-ins driven by the same promoter at the same genomic locus. We then recorded the mechanosensory responses from the gentle touch neuron ALM in strains carrying each variant from our toolkit. We also demonstrate the efficacy of these reagents in recording mechanosensory responses from the soma at low magnification and neural processes at high magnification. Finally, by making our library of reagents publicly available, we hope to facilitate the investigation of subcellular and multicellular calcium dynamics amongst the *C. elegans* research community.

## Materials and Methods

### Generation of GCaMP variants

We took an *in silico* approach to optimizing the coding sequence (CDS) of GCaMP6s (CHEN *et al*. 2013) for efficient expression in *C. elegans*. First, we created a nuclear-localized version of the fluorophore by adding the SV40 nuclear localization sequence (NLS) to its N-terminus and an *egl-13* NLS at the C-terminus (LYSSENKO *et al*. 2007). Next, we optimized this modified sequence for the worm genome using the *C. elegans* codon adapter (REDEMANN *et al*. 2011), which also inserted artificial introns into the CDS to enhance expression *in vivo*. The optimized sequence was then synthesized as a gene fragment (Integrated DNA Technologies) along with a fragment containing the *gpd-2/gpd-3* operonic linker (HUANG *et al*. 2001) with overlaps with the last exon of our nuclear-localized GCaMP6s and the first exon of our existing nuclear-localized mScarlet-I gene (STEVENSON *et al*. 2022). These two gene fragments were used directly for DNA assembly (New England Biolabs) into pDSP9, our mScarlet-I plasmid, to obtain pDSP23, a promoterless version of the bicistronic construct containing nuclear-localized GCaMP6s linked to nuclear-localized mScarlet-I.

The promoterless vectors for nuclear-localized GCaMP6f, 7s, and 7f variants were derived by site-directed mutagenesis (NEB Q5 Site-Directed Mutagenesis Kit) of the GCaMP6s sequence in pDSP23 to reproduce the mutations described for GCaMP6f (pDSP31) (CHEN *et al*. 2013) and jGCaMP7s pDSP32), 7f (pDSP33) (DANA *et al*. 2019). The cytosolic GCaMP6s, 6f, jGCaMP7s, and 7f variants (pHJL1-4, respectively) were derived from the nuclear-localized versions via PCR and DNA assembly to remove the SV40 and *egl-13* NLS. The membrane-localized GCaMP6s, 6f, jGCaMP7s, and 7f variants (pSiM1-4, respectively) were also derived from the nuclear-localized versions using the same methodology to remove the SV40 and *egl-13* NLS and add the C-terminal *ras-2* CAAX domain (CHEN *et al*. 2013; HISAMOTO *et al*. 2016) to both the GCaMP and mScarlet-I genes. All vectors were sequenced to verify correct assembly before further use. The primer sequences used for all constructions are provided in Table S2.

### Single-copy transgenesis of *C. elegans*

We created single-copy knock-ins of each of our constructs driven by the *mec-7* promoter, expressed in the gentle touch neurons (HAMELIN *et al*. 1992), using mos-mediated transgenesis (FRØKJÆR-JENSEN *et al*. 2012). The *mec-7* promoter was amplified from genomic DNA. This amplicon was assembled along with the bicistronic GCaMP-mScarlet-I construct of each variant into pDSP2, a kanamycin-resistant version of the pCFJ350 vector (FRØKJÆR-JENSEN *et al*. 2012). Oligo sequences for all PCRs are given in Table S2. Transgenic single-copy integrants were obtained following the MosSCI protocol (FRØKJÆR-JENSEN *et al*. 2012). Briefly, QL74, a gift from QueeLim Ch’ng, was used for all construct injections. Single copy integrants were screened using *peel-1* negative selection and loss of extrachromosomal co-injection markers. The resulting strains are listed in Table T1.

### Imaging/Data collection

All functional imaging experiments were performed on the same system using the same procedure described in Cho *et al*. (2017). Briefly, 2-day-old adult worms are loaded into and acclimated to the device and imaging light for 2 minutes. Baseline neural fluorescence was recorded for 10 seconds before delivering a 3-second 2.5 Hz mechanical stimulus. The neural activity following the mechanical stimulus was recorded for 60-90 seconds. All functional imaging was performed on a Leica DMIRB inverted microscope with a 40x air objective (N.A. 0.75) unless otherwise specified. Low magnification dynamics (Figure 3) were recorded using a 10x (N.A. 0.22) air objective. Video recordings were performed on a Hamamatsu EM-CCD with 100 ms exposure. Simultaneous two-color imaging was acquired with a DV2 beam splitter (Photometrics) and GFP/RFP filter set with the excitation coming from a projector source (STIRMAN *et al*. 2011).

### Analysis

Fluorescence intensities for each frame were obtained as previously described (CHO *et al*. 2017). Briefly, the GCaMP-only fluorescence and ratiometric, GCaMP plus RFP or mScarlet-I) values were computed using a customized MATLAB script. We primarily report ΔF (GCaMP only) values to minimize the effect of different red intensities in comparing GCaMP variants. The baseline fluorescence values were calculated using the frames prior to the stimulus.

## Results and Discussion

### Design of the transgenes

To provide a toolkit of GCaMPs tailored to *C. elegans* neuroscience research, we modified the most frequently used variants of GCaMP from mammalian studies. The *C. elegans*-specific toolkit features four variants with different response kinetics, each available with three different sub-cellular localizations. In addition to cytoplasmic fluorophores, we have created versions with localization to the nucleus and cell membrane to accommodate expansive imaging conditions. For example, the nuclear-localized expression may be helpful in imaging dense cell populations, such as in whole-brain imaging. The membrane-localized fluorophores can enable visualization of the calcium dynamics in the neurites. Further, we have incorporated two kinetic variants of GCaMP for different imaging needs: an *f* (fast) variant for faster rise and decay kinetics and an *s* (slow) variant for higher sensitivity. While the *s* variants typically offer higher signal-to-noise ratios, the *f* variants are useful in applications where responses to rapidly changing stimuli or interneuronal signaling must be observed.

To achieve the maximum possible signal-to-noise ratio, we first optimized the codon usage of GCaMP6s for the *C. elegans* genome. In addition, we also added three artificial introns into the coding sequence, as these are known to enhance transgene expression (OKKEMA *et al*. 1993). These modifications increase the number of *C. elegans* optimal codons from ∼27% in the original sequence to 85% while avoiding repetitive DNA sequences that could promote transgene silencing.

After creating our worm-optimized GCaMP6s, we used it as a template for site-directed mutagenesis to generate GCaMP6f (CHEN *et al*. 2013) and jGCaMP7s and f, once their sequences were publicly available (DANA *et al*. 2019). Many neural activity imaging experiments also use a red fluorophore, whose brightness is not linked to calcium levels, as a reference within each cell to help correct motion artifacts. To employ this strategy in our toolkit, we made our constructs bicistronic by including our codon-optimized mScarlet-I gene (STEVENSON *et al*. 2022) linked to the 3’ end of GCaMP gene using the *gpd-2/gpd-3* operonic linker sequence that promotes trans-splicing (HUANG *et al*. 2001). The bicistronic expression of GCaMP and mScarlet-I ensures similar protein levels within each cell, allowing the calcium-insensitive red fluorophore to be used for ratiometric normalization of the dynamic GCaMP signal and correct for motion artifacts.

### jGCaMP7 performs better than GCaMP6 *in vivo*

To compare and contrast the properties of *C. elegans*-optimized GCaMP6 and jGCaMP7 and to highlight the differences and utility of *in vivo* characteristics of the slow and fast variants, we expressed these single-copy transgenes in the mechanosensory neurons ALM, AVM, PLM, and PVM using the *mec-7* promotor (HAMELIN *et al*. 1992). We compare the *s* (slow) and *f* (fast) variants of GCaMP6 and jGCaMP7 because they have the maximum brightness and fastest decay dynamics, respectively. The ALM and AVM neurons are the sensory neurons responsible for responding to gentle touch in the anterior half of the body. To compare the functional activity of several GCaMP variants (6s, 7s, 6f, 7f), we used previously developed methods (CHO *et al*. 2017) to test the *in vivo* function of the reporters in ALM in response to consistent mechanical stimuli. In addition, we compared these GCaMP variants against an existing state-of-the-art GCaMP6m reporter strain, AQ3236: *ljSi2 [mec-7::GCaMP6m::SL2::TagRFP + unc-119(+)] II; unc-119(ed3) III*. (CHO *et al*. 2017). For this comparison, we use a standardized assay in which the stimulus is delivered from t=0s to t=3s. The magnitude of the stimulus is calibrated before each trial to maintain consistency between trials and devices. The imaging conditions, including camera setting and excitation power, were also standardized according to previously established methods (CHO *et al*. 2017). In order to compare the different localizations and GCaMP variants, we chose imaging conditions that allows for visualization of fluorophores without pixel saturation during recording of the brightest variants. The imaging conditions can be further optimized depending on which variant the user ultimately chooses. Under these stimulus and imaging conditions we find that the GCaMP variants behave as expected *in vivo* as previously reported *in vitro* (DANA *et al*. 2019).

The results of the previously described assay demonstrate that the new GCaMP reporter strains cover a range of properties such as peak brightness and decay time while performing as well or better than existing state-of-the-art reporters. Figure 1A shows heatmaps of the fluorescence change for many individuals carrying each reporter responding to the same mechanical stimuli. While all populations are shown to have some individuals with a large increase in fluorescence, the jGCaMP7s and 7f populations have much more activity on a per individual basis compared to their GCaMP6 counterparts. Figure 1B shows the average fluorescence for each population, normalized by their baseline brightness over time. The fluorescence increases from 0-3s, when the stimulus is applied, and then returns to the baseline brightness. The normalized peak brightness, representing dynamic range, is shown to be similar between all strains. However, when comparing the rate of decay, GCaMP6f is notably faster, and jGCaMP7s slower, than the other strains (Figure 1B). It is important to note that the amplitude of the peaks, while a fair comparison between strains, is not a representation of the brightest output these fluorophores can achieve in response to other stimuli. Mechanical stimulus of the ALM has been shown to produce around one fifth of the amplitude of a light response (NEKIMKEN *et al*. 2017). Furthermore, as we have used MoSCI to obtain single copy integrants, the copy number can be increased further to increase signal and to tune brightness to suit a particular use case.

**Figure 1:**
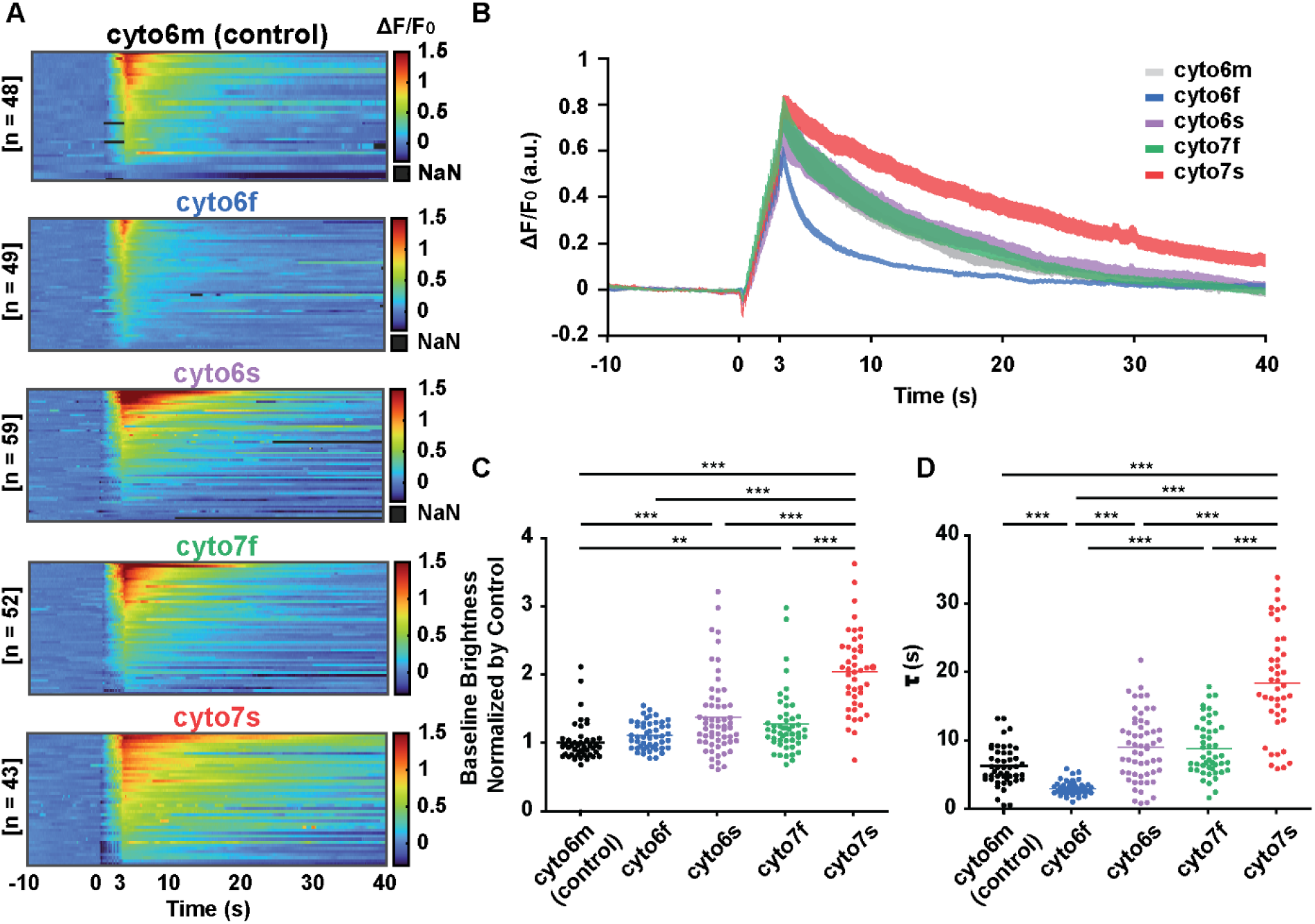
jGCaMP7 performs better than GCaMP6 *in vivo*. (A) Individual fluorescence intensity over time, with stimulus delivered at t = 0 s are shown for each developed cytosolic localized GCaMP strain as well as the existing AQ3236 strain which is used for comparison. B) GCaMP brightness, normalized by pre-stimulus baseline, increases then decreases in the mechanosensory neuron over time, where a controlled mechanical stimulus is applied from 0-3 seconds (dotted line). Amplitude of brightness relative to baseline and time to relax back to baseline vary depending on GCaMP variant. C) Baseline brightness of GCaMP of each developed cytosolic variant is compared to a previously developed GCaMP6m cytosolic reporter strain, normalizing all values by the average baseline brightness of the comparison strain. Notably, the newly developed 7f and 7s GCaMP variants are significantly brighter than the GCaMP6m strain. D) The time constants of the decays of each of the GCaMP variants are compared. Notably, 7s is found to decay at a significantly slower rate than all other strains and 6f is found to be significantly faster than all other variants. The Kruskal-Wallis test was performed, followed by Dunn’s multiple comparison test (** P < 0.01, *** P<0.001).

Comparing baseline brightness and decay rate of the different fluorophores, which are shown in Table 2, point to their potential for different experimental uses. When comparing baseline brightness, the jGCaMP7 variants perform significantly better than AQ3236 as well as the codon-optimized GCaMP6 strains (Figure 1C). All strains except the optimized GCaMP6f were significantly different in brightness compared to AQ3236, and jGCaMP7s is shown to be significantly brighter than all other fluorophores. As for rates of decay, the jGCaMP7s strain is shown to have the largest time constant, i.e. it is slowest to return to baseline brightness (Figure 1D), while GCaMP6f is shown to be significantly faster than the other variants.

**Table 2:** Brightness and decay speed of cytosolic GCaMP variants.

| GCaMP variant | Baseline Brightness normalized by 6m | $\tau$ (s) |
| --- | --- | --- |
| 6m | $1 \pm 0.28$ | $6.3 \pm 2.9$ |
| 6s | $1.45 \pm 0.57$ | $8.4 \pm 5.0$ |
| 7s | $2.05 \pm 0.60$ | $18.0 \pm 8.1$ |
| 6f | $1.11 \pm 0.20$ | $2.7 \pm 2.0$ |
| 7f | $1.28 \pm 0.46$ | $8.5 \pm 4.1$ |

For example, the optimized jGCaMP7s has the highest baseline brightness (260% increase from unoptimized 6m), but also the slowest kinetics (t_1/2, fall_ = 18 ± 8 s). The optimized GCaMP6f has the lowest baseline brightness (11% increase from unoptimized 6m), but the fastest kinetics (t_1/2, fall_ = 2.7 ± 2.0 s).

The high baseline brightness of jGCaMP7s combined with its slow decay time makes it best suited for situations where it is not required for the fluorophore to decay quickly, or in fact if a slow decay rate is desirable. It is also useful in situations where the highest possible fluorescence intensity is required, such as imaging at low magnification or lower excitation power. On the other hand, jGCaMP7f is shown to be as fast as the control strain while also being brighter, which will allow experiments to be performed at higher camera speeds. This indicates that jGCaMP7f will be useful when both high temporal resolution and signal are required. Its peak brightness is significantly greater than that of jGCaMP6f, and decay rate is about 2 times faster than jGCaMP7s (Figure 1B, D). These properties are useful for measuring neuronal activity at faster time scales, such as responses to a fast, time-varying stimulus. GCaMP6f has the fastest decay rate but has a tradeoff of low dynamic range, which may prevent its use in measurements of low magnitude stimuli.

### Subcellular localization provides a toolkit for multiple applications

To compare the GCaMP localization, we used the same neurons and measured the baseline and excited fluorescence (Figure 2A). We see that GCaMP localized to the nucleus could be best used for multicell imaging of a cluster of neurons that would be otherwise unresolvable, e.g. as in imaging chemosensory head neuron activities. Membrane bound GCaMP may allow for better study of compartmentalized or subcellular calcium signaling, e.g. in neurites. For example, it could be used for examining local sensory input, e.g. in proprioception (TAO *et al*. 2019).

**Figure 2:**
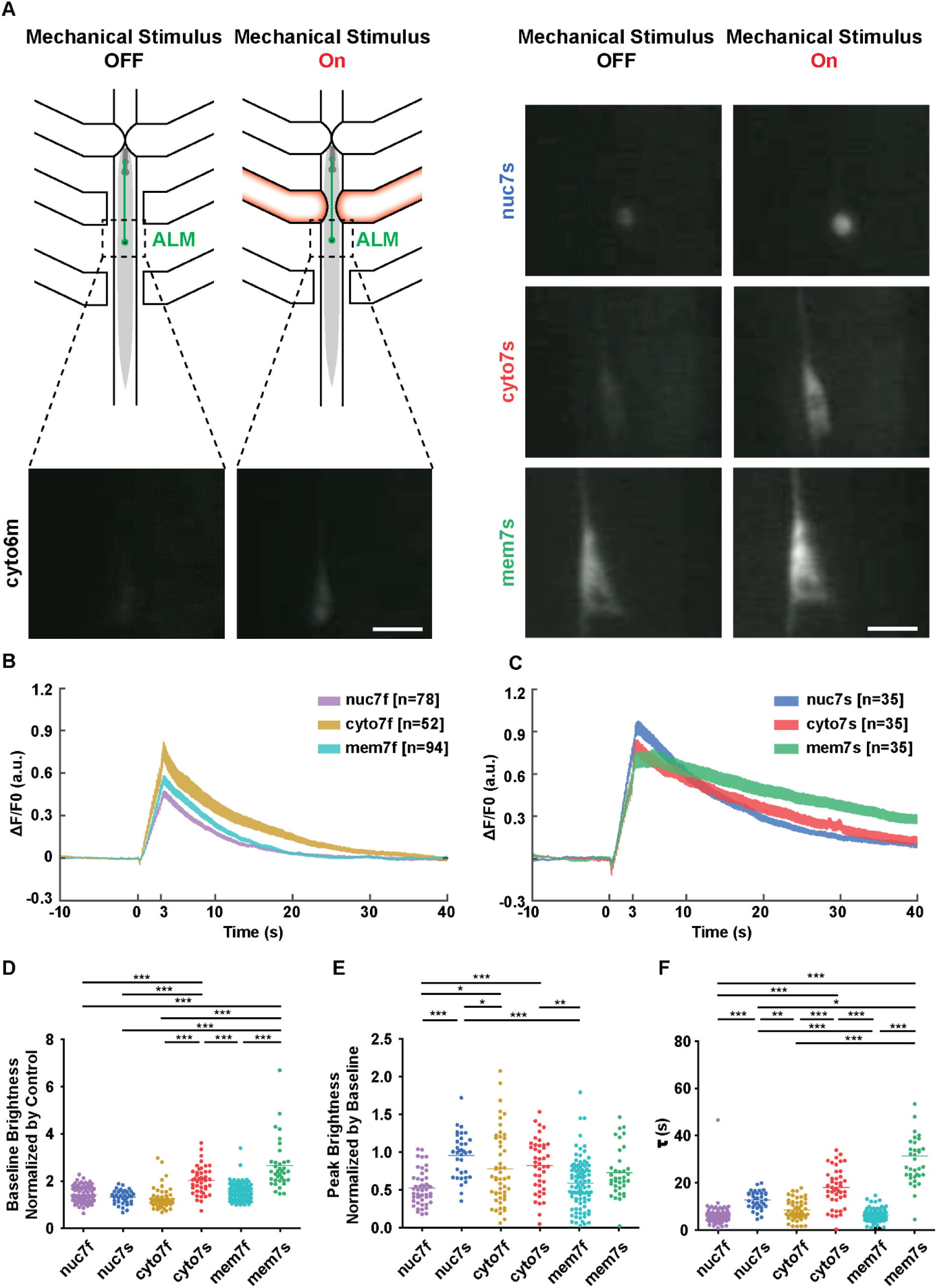
Quantitative comparison between different subcellular localizations. A) Mechanical stimulus is delivered via pneumatic valves, leading to an increase in neuronal fluorescence. Example baseline (left) and excited (right) fluorescent images are shown for GCaMP6m comparison strain, and for the cytosolic, nuclear, and membrane subcellular localizations for jGCaMP7s. Scale bar: 3μm. B) Average change in fluorescence over time normalized by baseline activity, with stimulus occurring at t=0 s, for all three localizations of jGCaMP7f and 7s (C). The measured brightness increases after stimulus delivery before reaching a peak and returning to baseline. Different localizations demonstrate small variations in various parameters such as decay time and peak height. D) Comparison of baseline brightness normalized by control strain for all subcellular localizations of jGCaMP7f and jGCaMP7s. All strains are found to be, on average, brighter than control strain. The nuclear localized strains are found to be dimmest while membrane bound jGCaMP7s is found to be the brightest. E) Peak GCaMP brightness normalized by baseline brightness for each subcellular localization of jGCaMP7f and 7s. Generally, the jGCaMP7f variants are seen to have somewhat lower dynamic range. F) Comparison of speed of decay for each subcellular localization. All localizations of the jGCaMP7f variants are found to be significantly faster than all of the 7s variants, with membrane 7s decaying at the slowest rate and membrane 7f with the most consistently fast decay. The Kruskal-Wallis test was performed, followed by Dunn’s multiple comparison test (* P < 0.05, ** P < 0.01, *** P<0.001).

We next characterize and compare different subcellular localizations by imaging their functional response using the same standardized assay described above (Figure 2 B-F). We again use an existing state-of-the-art reporter strain, AQ3236 (CHO *et al*. 2017) as a control for comparison with the optimized variants. By examining baseline brightness normalized by average brightness of the control strain (Figure 2D), we observed that all new jGCaMP7 variants are brighter than the previously used control strain. The membrane jGCaMP7s is the brightest, with some extreme outliers at over six times brighter than the control. The rest of the strains are closer in brightness with general trends of jGCaMP7s being brighter than jGCaMP7f regardless of localization, and the membrane and cytosolic localization being brighter than the nuclear localized strain. Dynamic range is demonstrated through measuring neuronal response to a stimulus delivery normalized by the baseline brightness (Figure 2E). Again, the jGCaMP7s strains are shown to have a higher dynamic range than their jGCaMP7f counterparts for each localization. Nuclear localized jGCaMP7f demonstrated the worst range at around a 50% increase from baseline brightness, whereas the nuclear and cytosolic localized jGCaMP7s variants demonstrate around a 100% increase from baseline. The membrane bound strains demonstrate slightly lower dynamic range comparatively. The decay dynamics are also characterized (Figure 2F), and, as expected, the jGCaMP7s variants are slower to decay compared to their jGCaMP7f counterparts. Notably membrane jGCaMP7s is found to take by far the longest to decay from excited brightness whereas all three jGCaMP7f localizations decay at a rapid rate.

These results demonstrate that although nuclear localization decreases the imaging area for neurons and can be beneficial when imaging highly clustered neurons, it also decreases baseline brightness. In contrast, membrane localization increases the baseline brightness. This is beneficial when imaging at lower magnification; further, membrane localization would allow small structures such as dendrites to be imaged. It additionally makes this localization best suited for systems requiring neuron tracking as the higher baseline brightness reduces the chances of the ROI being lost or confused with, for example, autofluorescence, if a somewhat lower signal-to-noise ratio can be tolerated.

### New GCaMP variants allow for novel opportunities in functional imaging

Functional imaging at low magnification opens up new opportunities to, for example, record more accurate neuronal activity during behavioral tracking, or image the neural activity of many animals simultaneously using a single objective during a population-wide stimulus, such as a chemical cue. Our newly developed reporter strains allow lower magnification imaging because of their increased brightness. We demonstrate this by recording the fluorescence of the ALM neuron tagged with membrane localized jGCaMP7s while it is undergoing the same stimulus regime described above, using a lens with much lower magnification and NA (10x, N.A. 0.22) compared to the objective used in the previously described assay (40x, N.A. 0.75) (Figure 3A). The baseline fluorescence of the gentle touch neurons is bright enough to easily identify through the eyepiece by eye, and it allows for imaging neurons in freely moving worms (Figure 3B). This also facilitates proper identification and adjusting the optical focus for specific neurons at low magnification. We further demonstrate functional imaging in a mechanically stimulated neuron is possible at this magnification, which has not been shown before. The brighter membrane 7s fluorophore allows observation of a significant increase in fluorescence (Figure 3C) detectable even at low magnification, with a low NA (0.22) air objective and using relatively low excitation from a LCD projector source (STIRMAN *et al*. 2011) as opposed to a laser. In comparison, 7f or AQ3236 are too dim to image under the same condition.

**Figure 3:**
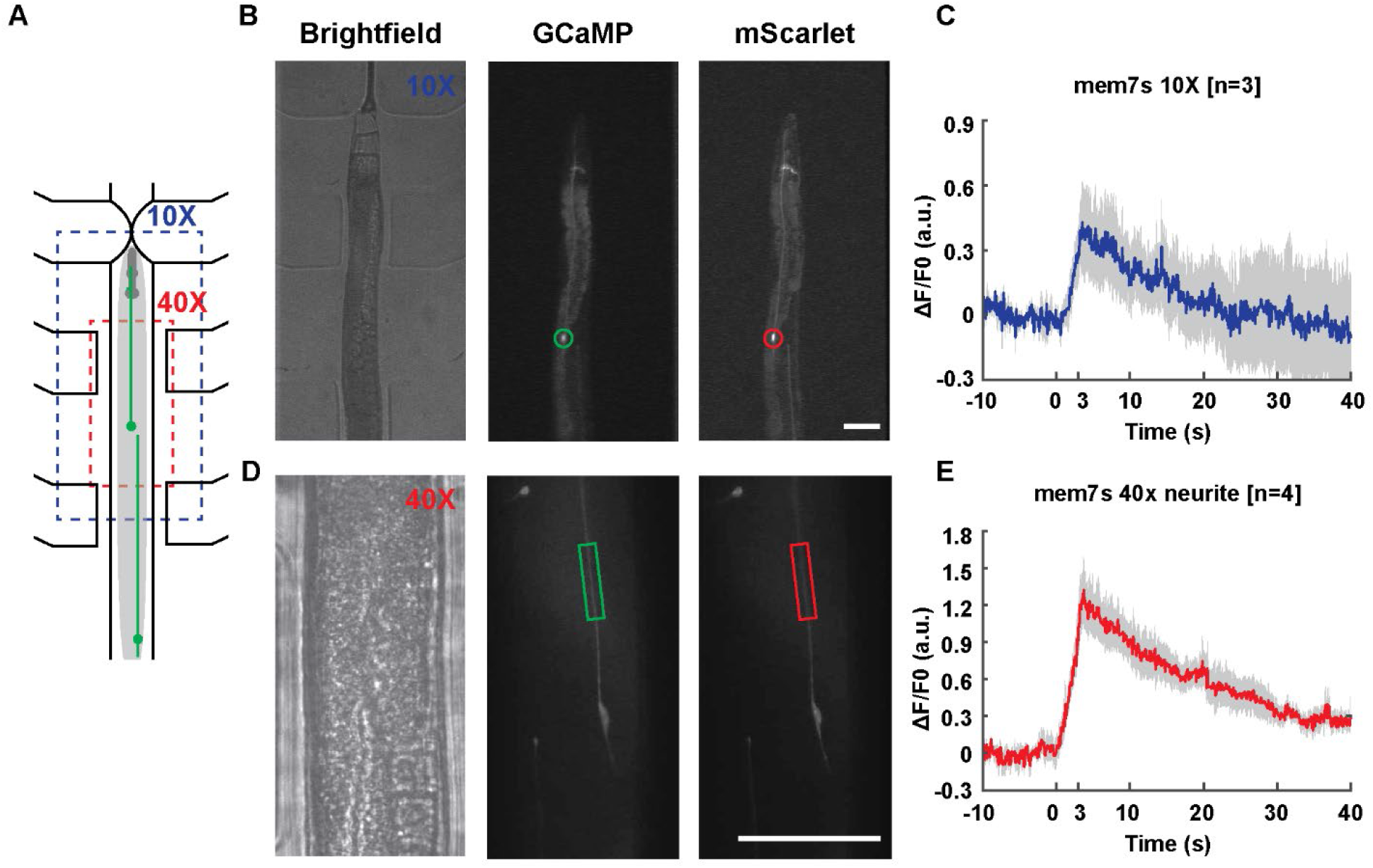
Low magnification and neurite functional imaging are demonstrated using newly developed strain. A) Diagram demonstrating increasing field of view at lower 10x magnification compared to the 40x magnification used in previous figures. B) Brightfield (left), and fluorescent GCaMP (middle), and mScarlet (right) images taken at lower magnification, with nerve ring and ALM neuron body visible. Green and red circles indicate ROI for calculation of fluorescent intensity. Scale bar: 50 μm C) Functional imaging is demonstrated to still be possible at 10x magnification using our microscopy platform and the newly developed membrane localized GCaMP7s reporter strain, with a significant increase in normalized fluorescence compared to baseline activity. D) Brightfield (left), and fluorescent GCaMP (middle), and mScarlet (right) images taken at standard magnification demonstrate that the neuron process is clearly visible. Green and red boxes indicate ROI for calculation of fluorescent intensity. Scale bar: 50 μm E) Functional imaging of neurite is demonstrated to be possible for the membrane jGCaMP7s strain under these imaging conditions.

The subcellular localization of GCaMP also allows for investigation of intracellular processing or local cellular response. Membrane bound GCaMP allows us to look at how information is passed from the neurite to the cell body. Our membrane bound GCaMP strain has high enough signal-to-noise ratio in the neurite of the mechanosensory neurons to quantify calcium transients in the neurite. We demonstrate this by analyzing the neurite’s response to mechanical stimulus in a region of the neurite that is 60 μm to 100 μm (150-250 pixels) anterior to the cell body (Figure 3E). While this portion of the process is not visible in the cytosolic and nuclear localizations, we can extract robust GCaMP responses in the membrane-bound localization, based on segmentation of the red channel. Using the same procedure as previously described, we see that the half-life of the signal in the neurite is 16.4 s (Figure 3E). This is approximately half of the 31 s half-life observed for the cell body, shown in Figure 2F. We speculate that the difference in decay time between neurite and cell body could indicate that the cell body is integrating the mechanosensory signal from the neurite.

In summary, this work provides a comparison of GCaMP6s, 6f, and jGCaMP7s, 7f variants with different subcellular localization in *C. elegans*. We showed both jGCaMP7 variants are brighter than the GCaMP6 counterparts. *In vivo* results showed that the decay time of 6f is faster or similar to 7f when there is a large signal (greater than 40 action potentials), indicating that for *C. elegans*, which has few spiking neurons, 6f may show more favorable decay rates. However, jGCaMP7f has greater signal (ΔF/F_0_) compared to GCaMP6f, so there seems to be a tradeoff between signal and decay rate in *C. elegans*. Among the slow variants (6s and 7s), a similar tradeoff exists, with jGCaMP7s being slower but brighter and GCaMP6s being faster with less signal. However, GCaMP6s has similar decay kinetics to jGCaMP7f, but jGCaMP7f is brighter, so there seems to be little advantage to using GCaMP6s compared to the other three variants that we compared.

We also found that membrane localization provides greater signal in *C. elegans*. Nuclear localization of the protein allows for neurons in a dense population to be distinguished, but our results indicate that this decreases the signal. While the ΔF/F_0_ was not significantly different between localization, the effect of localization on size of the neuron and raw fluorescence values means that the overall signal is higher in membrane localized variants. This enhanced signal allowed us to record calcium activity even in a low magnification low NA condition. The ability to image neural activity in freely moving worms has often required custom built microscopes in order to get the magnification necessary to record the low signal. In sparsely labeled neurons, it would be advantageous to use membrane-bound GCaMP, as the greater signal allows for lower magnification to be used. As new GCaMP variants continue to be developed and improved, the plasmids provided by our toolkit can be used as a foundation to produce additional *C. elegans* optimized plasmids and be further mutated to a desired new GCaMP coding sequence. For example, the GENIE group at the HHMI Janelia research campus has recently developed their next generation jGCaMP8 series of reporters (ZHANG *et al*. 2021). These reporters have been developed for *Drosophila* and mammalian studies. However, our codon-optimized toolkit can easily be modified by the *C. elegans* research community to covert our GCaMP6s into any of the new jGCaMP8 variants using site-directed mutagenesis. This work demonstrates the use of codon optimization to improve expression, and brightness, in *C. elegans*. Other model organisms utilizing GCaMP may also benefit from codon optimizing their GCaMP of choice, as it will likely improve performance and brightness.

## Data availability

Strains and plasmids are available upon request. Individual functional response data are available upon request. Code used for image processing and data analysis is available upon request.

## Acknowledgements

The authors wish to acknowledge Dr. QueeLim Ch’ng for providing QL74 strain all MosSCI construct injections. This work is partially supported by funding from the National Institutes of Health (R01AG056436, R01NS115484, and R01NS096581 to HL) and the National Science Foundation (1764406 and 1707401 to HL).

## Supporting Information

**Table T1:** Transgenic strains.

| Strain Name | GCaMP variant | Subcellular localization | Genotype | Reference |
| --- | --- | --- | --- | --- |
| AQ3236 | 6m | Cytosolic | *ljSi2 [mec-7::GCaMP6m::SL2::TagRFP + unc-119(+)] II; unc-119(ed3) III.* | (CHO *et al.* 2017) |
| GT349 | 6s | Membrane | *aSi25[lox2272 Cbr-unc-119(+) lox2272 + mec-7p::GCaMP6s::ras-2CAAX::SL2::mScarlet-I::ras-2CAAX] II; unc-119(ed3) III.* | This work |
| GT350 | 7s | Membrane | *aSi26[lox2272 Cbr-unc-119(+) lox2272 + mec-7p::GCaMP7s::ras-2CAAX::SL2::mScarlet-I::ras-2CAAX] II; unc-119(ed3) III.* | This work |
| GT370 | 6f | Membrane | *aSi29[lox2272 Cbr-unc-119(+) lox2272 + mec-7p::GCaMP6f::ras-2CAAX::SL2::mScarlet-I::ras-2CAAX] II; unc-119(ed3) III.* | This work |
| GT372 | 7f | Membrane | *aSi31[lox2272 Cbr-unc-119(+) lox2272 + mec-7p::GCaMP7f::ras-2CAAX::SL2::mScarlet-I::ras-2CAAX] II; unc-119(ed3) III.* | This work |
| GT324 | 6s | Nuclear Localized | *aSi4[lox2272 Cbr-unc-119(+) lox2272 + mec-7p::NLS::GCaMP6s::egl-13NLS::SL2::NLS::mScarlet-I::egl-13NLS] II; unc-119(ed3) III.* | This work |
| GT330 | 7s | Nuclear Localized | *aSi8[lox2272 Cbr-unc-119(+) lox2272 + mec-7p::NLS::GCaMP7s::egl-13NLS::SL2::NLS::mScarlet-I::egl-13NLS] II; unc-119(ed3) III.* | This work |
| GT346 | 6f | Nuclear Localized | *aSi22[lox2272 Cbr-unc-119(+) lox2272 + mec-7p::NLS::GCaMP6f::egl-13NLS::SL2::NLS::mScarlet-I::egl-13NLS] II; unc-119(ed3) III.* | This work |
| GT347 | 7f | Nuclear Localized | *aSi23[lox2272 Cbr-unc-119(+) lox2272 + mec-7p::NLS::GCaMP7f::egl-13NLS::SL2::NLS::mScarlet-I::egl-13NLS] II; unc-119(ed3) III.* | This work |
| GT374 | 6s | Cytosolic | *aSi33[lox2272 Cbr-unc-119(+) lox2272 + mec7p::GCaMP6s::SL2::mScarlet-I] II; unc-119(ed3) III.* | This work |
| GT375 | 7s | Cytosolic | *aSi27[lox2272 Cbr-unc-119(+)*<br>*lox2272 + mec-7p::GCaMP7s::SL2::mScarlet-I]*<br>*II; unc-119(ed3) III.* | This work |
| GT376 | 6f | Cytosolic | *aSi35[lox2272 Cbr-unc-119(+)*<br>*lox2272 + mec7-p::GCaMP6f::SL2::mScarlet-I]*<br>*II; unc-119(ed3) III.* | This work |
| GT377 | 7f | Cytosolic | *aSi36[lox2272 Cbr-unc-119(+)*<br>*lox2272 + mec7-p::GCaMP7f::SL2::mScarlet-I]*<br>*II; unc-119(ed3) III.* | This work |

**Figure S1.**
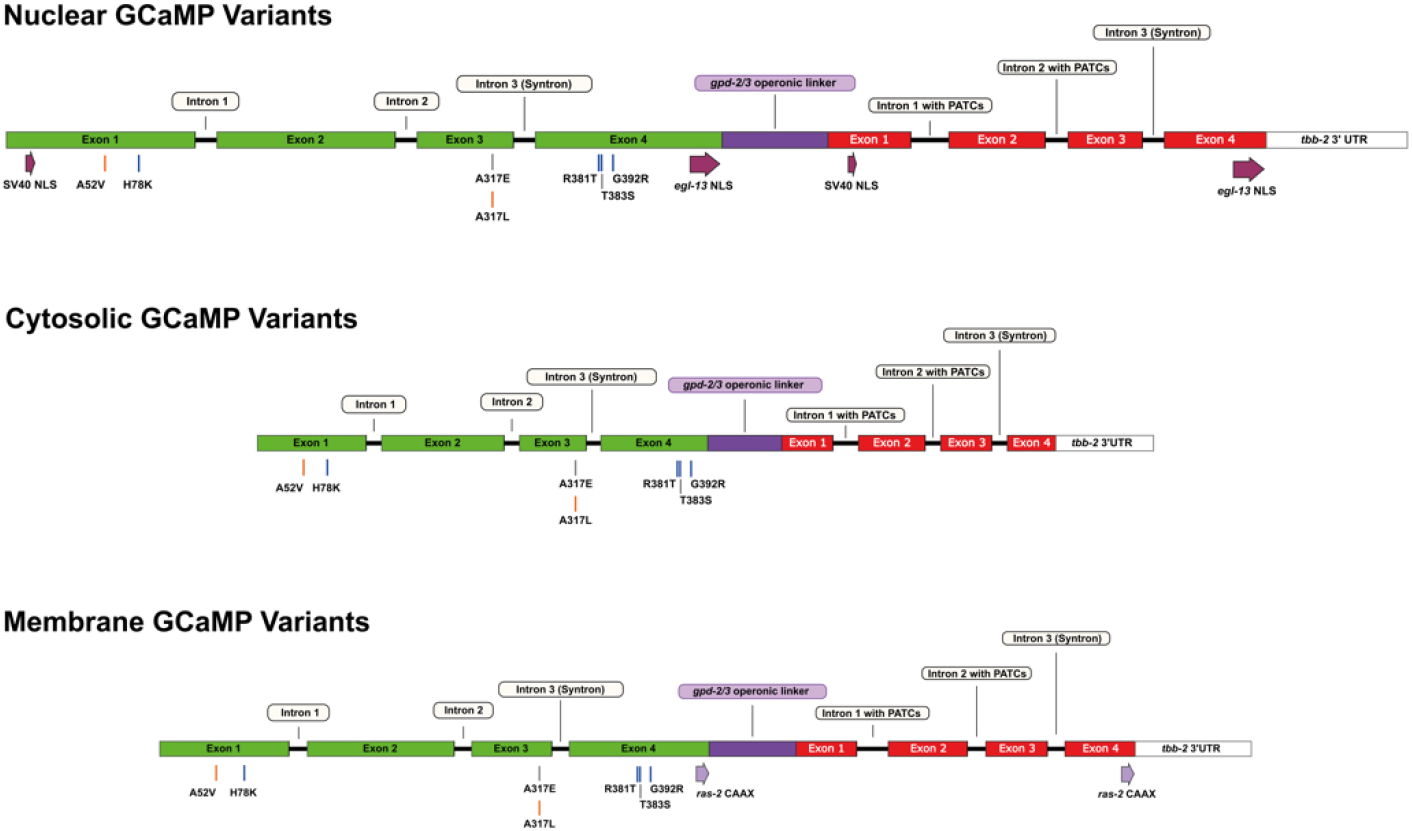
*C. elegans*-optimized GCaMP reporter toolkit. Schematic representation of the codon-optimized bicistronic GCaMP and mScarlet-I reporter genes. The exons for the GCaMP and mScarlet-I are color-coded in green and red, respectively. Amino acid substitutions resulting in the 6f (grey), 7s (orange), and 7f (blue) variants from GCaMP6s are indicated below the schematic for each variant. For convenience, we use the numbering scheme used in Dana et al, (2019). (Top) Nuclear-localized variants utilize two NLS sequences in each fluorophore, with the SV40 and *egl-13* NLS encoded at the 5’ and 3’ ends of the coding sequence, respectively. (Middle) Cytosolic variants were created by removing the NLS sequences from their nuclear-localized counterparts (see Methods). (Bottom) Membrane-localized variants were created from their cytosolic counterparts through the addition of a CAAX domain from the *ras-2* gene (see Methods).

## References

Ahrens, M. B., M. B. Orger, D. N. Robson, J. M. Li and P. J. Keller, 2013 Whole-brain functional imaging at cellular resolution using light-sheet microscopy. Nature Methods 10: 413–420.

Bindels, D. S., L. Haarbosch, L. van Weeren, M. Postma, K. E. Wiese et al., 2017 mScarlet: a bright monomeric red fluorescent protein for cellular imaging. Nat Methods 14: 53–56.

Chalasani, S. H., N. Chronis, M. Tsunozaki, J. M. Gray, D. Ramot et al., 2007 Dissecting a circuit for olfactory behaviour in Caenorhabditis elegans. Nature 450: 63 70.

Chen, T.-W., T. J. Wardill, Y. Sun, S. R. Pulver, S. L. Renninger et al., 2013 Ultrasensitive fluorescent proteins for imaging neuronal activity. Nature 499: 295-300.

Cho, Y., D. A. Porto, H. Hwang, L. J. Grundy, W. R. Schafer et al., 2017 Automated and controlled mechanical stimulation and functional imaging in vivo in C. elegans. Lab on a Chip 17: 2609–2618.

Clapham, D. E., 2007 Calcium signaling. Cell 131: 1047–1058.

Dana, H., Y. Sun, B. Mohar, B. K. Hulse, A. M. Kerlin et al., 2019 High-performance calcium sensors for imaging activity in neuronal populations and microcompartments. Nature methods 16: 649–657.

Flavell, S. W., N. Pokala, E. Z. Macosko, D. R. Albrecht, J. Larsch et al., 2013 Serotonin and the Neuropeptide PDF Initiate and Extend Opposing Behavioral States in C. elegans. Cell 154: 1023 1035.

Frøkjær-Jensen, C., M. W. Davis, M. Ailion and E. M. Jorgensen, 2012 Improved Mos1-mediated transgenesis in C. elegans. Nat Methods 9: 117–118.

Gordus, A., N. Pokala, S. Levy, S. W. Flavell and C. I. Bargmann, 2015 Feedback from network states generates variability in a probabilistic olfactory circuit. Cell 161: 215-227.

Ha, H.-i., M. Hendricks, Y. Shen, C. V. Gabel, C. Fang-Yen et al., 2010 Functional Organization of a Neural Network for Aversive Olfactory Learning in Caenorhabditis elegans. Neuron 68: 1173–1186.

Hamelin, M., I. M. Scott, J. C. Way and J. G. Culotti, 1992 The mec-7 beta-tubulin gene of Caenorhabditis elegans is expressed primarily in the touch receptor neurons. EMBO J 11: 2885–2893.

Hendricks, M., H. Ha, N. Maffey and Y. Zhang, 2012 Compartmentalized calcium dynamics in a C. elegans interneuron encode head movement. Nature 487: 99–103.

Hisamoto, N., Y. Nagamori, T. Shimizu, S. I. Pastuhov and K. Matsumoto, 2016 The C. elegans Discoidin Domain Receptor DDR-2 Modulates the Met-like RTK–JNK Signaling Pathway in Axon Regeneration. PLOS Genetics 12: e1006475.

Hong, M., L. Ryu, M. C. Ow, J. Kim, A. R. Je et al., 2017 Early Pheromone Experience Modifies a Synaptic Activity to Influence Adult Pheromone Responses of C. elegans. Current Biology 27: 3168-3177.e3163.

Huang, T., S. Kuersten, A. M. Deshpande, J. Spieth, M. MacMorris et al., 2001 Intercistronic region required for polycistronic pre-mRNA processing in Caenorhabditis elegans. Mol Cell Biol 21: 1111–1120.

Kato, S., Harris S. Kaplan, T. Schrödel, S. Skora, Theodore H. Lindsay et al., 2015 Global Brain Dynamics Embed the Motor Command Sequence of Caenorhabditis elegans. Cell 163: 656–669.

Kim, D. H., J. Kim, J. C. Marques, A. Grama, D. G. C. Hildebrand et al., 2017 Pan-neuronal calcium imaging with cellular resolution in freely swimming zebrafish. Nature Methods 14: 1107–1114.

Li, Y., A. Mathis, B. F. Grewe, J. A. Osterhout, B. Ahanonu et al., 2017 Neuronal Representation of Social Information in the Medial Amygdala of Awake Behaving Mice. Cell 171: 1176-1190.e1117.

Liu, Q., P. B. Kidd, M. Dobosiewicz and C. I. Bargmann, 2018 C. elegans AWA Olfactory Neurons Fire Calcium-Mediated All-or-None Action Potentials. Cell 175: 57-70.e17.

Lyssenko, N. N., W. Hanna-Rose and R. A. Schlegel, 2007 Cognate putative nuclear localization signal effects strong nuclear localization of a GFP reporter and facilitates gene expression studies in Caenorhabditis elegans. Biotechniques 43: 596, 598, 560.

Nekimken, A. L., H. Fehlauer, A. A. Kim, S. N. Manosalvas-Kjono, P. Ladpli et al., 2017 Pneumatic stimulation of C. elegans mechanoreceptor neurons in a microfluidic trap. Lab on a Chip 17: 1116–1127.

Nguyen, J. P., F. B. Shipley, A. N. Linder, G. S. Plummer, M. Liu et al., 2016 Whole-brain calcium imaging with cellular resolution in freely behaving Caenorhabditis elegans. Proceedings of the National Academy of Sciences 113: E1074–E1081.

Okkema, P. G., S. W. Harrison, V. Plunger, A. Aryana and A. Fire, 1993 Sequence requirements for myosin gene expression and regulation in Caenorhabditis elegans. Genetics 135: 385–404.

Parker, N. F., C. M. Cameron, J. P. Taliaferro, J. Lee, J. Y. Choi et al., 2016 Reward and choice encoding in terminals of midbrain dopamine neurons depends on striatal target. Nature Neuroscience 19: 845–854.

Redemann, S., S. Schloissnig, S. Ernst, A. Pozniakowsky, S. Ayloo et al., 2011 Codon adaptation-based control of protein expression in C. elegans. Nat Methods 8: 250–252.

Stevenson, Z. C., M. J. Moerdyk-Schauwecker, S. A. Banse, D. S. Patel, H. Lu et al., 2022.

Stirman, J. N., M. M. Crane, S. J. Husson, S. Wabnig, C. Schultheis et al., 2011 Real-time multimodal optical control of neurons and muscles in freely behaving Caenorhabditis elegans. Nature Methods 8: 153–158.

Tao, L., D. Porto, Z. Li, S. Fechner, S. A. Lee et al., 2019 Parallel Processing of Two Mechanosensory Modalities by a Single Neuron in C. elegans. Developmental Cell 51: 617-631.e613.

Venkatachalam, V., N. Ji, X. Wang, C. Clark, J. K. Mitchell et al., 2016 Pan-neuronal imaging in roaming Caenorhabditis elegans. Proceedings of the National Academy of Sciences 113: E1082–E1088.

Wirak, G. S., J. Florman, M. J. Alkema, C. W. Connor and C. V. Gabel, 2022 Age-associated changes to neuronal dynamics involve a disruption of excitatory/inhibitory balance in C. elegans. Elife 11.

Wu, T., M. Ge, M. Wu, F. Duan, J. Liang et al., 2023 Pathogenic bacteria modulate pheromone response to promote mating. Nature 613: 324–331.

Zaslaver, A., I. Liani, O. Shtangel, S. Ginzburg, L. Yee et al., 2015 Hierarchical sparse coding in the sensory system of Caenorhabditis elegans. Proceedings of the National Academy of Sciences 112: 1185–1189.

Zhang, Y., M. Rózsa, Y. Liang, D. Bushey, Z. Wei et al., 2021 Fast and sensitive GCaMP calcium indicators for imaging neural populations. bioRxiv: 2021.2011.2008.467793.

Zong, W., H. A. Obenhaus, E. R. Skytøen, H. Eneqvist, N. L. d. Jong et al., 2022 Large-scale two-photon calcium imaging in freely moving mice. Cell 185: 1240-1256.e1230.

